# Forward genetic screen of human transposase genomic rearrangements

**DOI:** 10.1101/061770

**Authors:** Anton G. Henssen, Eileen Jiang, Jiali Zhuang, Luca Pinello, Nicholas D. Socci, Richard Koche, Mithat Gonen, Camila M. Villasante, Scott A. Armstrong, Daniel E. Bauer, Zhiping Weng, Alex Kentsis

## Abstract

**Background.** Numerous human genes encode potentially active DNA transposases or recombinases, but our understanding of their functions remains limited due to shortage of methods to profile their activities on endogenous genomic substrates. **Results.** To enable functional analysis of human transposase-derived genes, we combined forward chemical genetic hypoxanthine-guanine phosphoribosyltransferase 1 (*HPRT1*) screening with massively parallel paired-end DNA sequencing and structural variant genome assembly and analysis. Here, we report the *HPRT1* mutational spectrum induced by the human transposase PGBD5, including PGBD5-specific signal sequences (PSS) that serve as potential genomic rearrangement substrates. **Conclusions.** The discovered PSS motifs and high-throughput forward chemical genomic screening approach should prove useful for the elucidation of endogenous genome remodeling activities of PGBD5 and other domesticated human DNA transposases and recombinases.

## Background

The human genome contains over twenty genes with similarity to DNA transposases (1). In addition, transposons are a major source of structural genetic variation in human populations (2). Recently, human THAP9 and PGBD5 have been found to mobilize transposons in human cells (3, 4). This discovery raises the possibility that, similar to the RAG1 recombinase (5), these endogenous human transposases may catalyze human genome rearrangements during normal somatic cell development or in distinct disease states. The human genome contains thousands of genetic elements with apparent sequence similarity to transposons, but their evolutionary divergence hinders the identification of elements that may serve as substrates for endogenous human transposases in general (6), and PGBD5 in particular (4).

In classical genetics, forward chemical genetic screens have been successfully used to identify spontaneous mutations in bacteria, yeast and fly (7–10). Such approaches use DNA sequencing of cells based on chemical resistance due to positive or negative phenotypic selection. For forward genetics of mammalian and human cells, mutational analysis of the hypoxanthine-guanine phosphoribosyltransferase 1 (*HPRT1*) gene based on the resistance to toxic purine analogues such as 8-aza- or 6-thio-guanine (referred to as thioguanine) has been used; for overview, see (11). Analysis of *HPRT1* has several advantages for forward genetic screens: i) *HPRT1* is on the X chromosome and therefore functionally hemizygous, ii) *HPRT1* encodes a single domain globular protein in which alterations of any of its nine exons are expected to affect enzymatic activity, and iii) mutations can be selected both positively and negatively, enabling the specific identification of distinct mutations, as opposed to general factors controlling cellular genomic stability. Indeed, HPRT1-based forward genetic screens have been successfully used to characterize chemical mutagens (12, 13). In human lymphocytes, this assay has also been used to identify RAG1-mediated mutations of *HPRT1*, and to elucidate cryptic recombination signal sequences (14, 15).

Here, we sought to develop a forward genetic screening approach suitable for the elucidation of endogenous genomic substrates of human DNA transposases and recombinases. Depending on cell type and presence of endogenous co-factors, this assay should allow for DNA transposition and recombination, or alternatively, nuclease-mediated DNA rearrangements facilitated by endogenous DNA sequence substrate preferences. Using negative and positive thioguanine resistance selection, combined with massively parallel DNA sequencing, we used *HPRT1* screening to investigate the nuclease activity of PGBD5 on human genomic substrates.

## Results

The human *HPRT1* gene contains 12 annotated DNA transposon copies (Supplementary Table 1). These transposons are only distantly related to *piggyBac* transposons that are evolutionarily related to the potential substrates of PGBD5 (4). We hypothesized that under strong selective pressure, PGBD5 may exhibit enzymatic activity on sequences in the *HPRT1* gene with sufficient similarity to its endogenous substrates, be they *piggyBac-related* sequences or not. To test this hypothesis, we adapted the *HPRT1* mutation assay in which cells containing inactivating *HPRT1* mutations can be negatively or positively selected by growth in media containing hypoxanthine-aminopterin-thymidine (HAT) or thioguanine, respectively (16). To maximize the sensitivity of this assay, we used male BJ fibroblasts containing a single copy of the X-linked *HPRT1* gene (17).

To analyze specific changes induced by human PGBD5, we generated isogenic cell lines by using lentiviral transduction to express GFP-PGBD5 and control GFP, and confirmed stable and equal transgene expression using immunoblotting (Fig. 1A). To eliminate *HPRT1* variants induced by spontaneous gene mutations and enable the specific selection of those induced by PGBD5, we grew cells in the presence of HAT medium for 15 doublings (18). We confirmed that the expression of GFP-PGBD5 did not alter the intrinsic sensitivity of BJ cells to thioguanine, as assessed using clonogenic assays and analysis of the dose-response to thioguanine by these cells (Fig. 1B and Fig. 1C). The clonogenic efficiencies of thioguanine resistance induction were 0.027 and 0.028 for cells expressing GFP and GFP-PGBD5, respectively, estimated from the number of thioguanine-resistant colonies, consistent with the lack of general mutagenic activity by PGBD5 (12, 13). To generate cells with *de novo HPRT1* mutations, we grew cells expressing GFP-PGBD5 or GFP control in the presence of thioguanine for 10 cell divisions, corresponding to approximately one month in culture. To confirm the generation of clones with inactivating *HPRT1* mutations, we determined the ability of thioguanine-selected cells to grow in the HAT medium. We observed more than 3.5-fold decrease in the number of viable cells upon HAT treatment, indicating that the majority of the cells had acquired resistance to thioguanine (Fig. 1D). Consistent with the inactivation of *HPRT1* in these cells, thioguanine-selected cells exhibited increased resistance to acute treatment with thioguanine as compared to unselected control cells (Fig. 1E). In agreement with this notion, we observed no detectable HPRT1 protein expression by immunoblotting (Fig. 1F), suggesting that the majority of thioguanine-resistant cells inactivated *HPRT1* due to mutations that led to the loss of HPRT1 protein expression.

**Figure 1.**
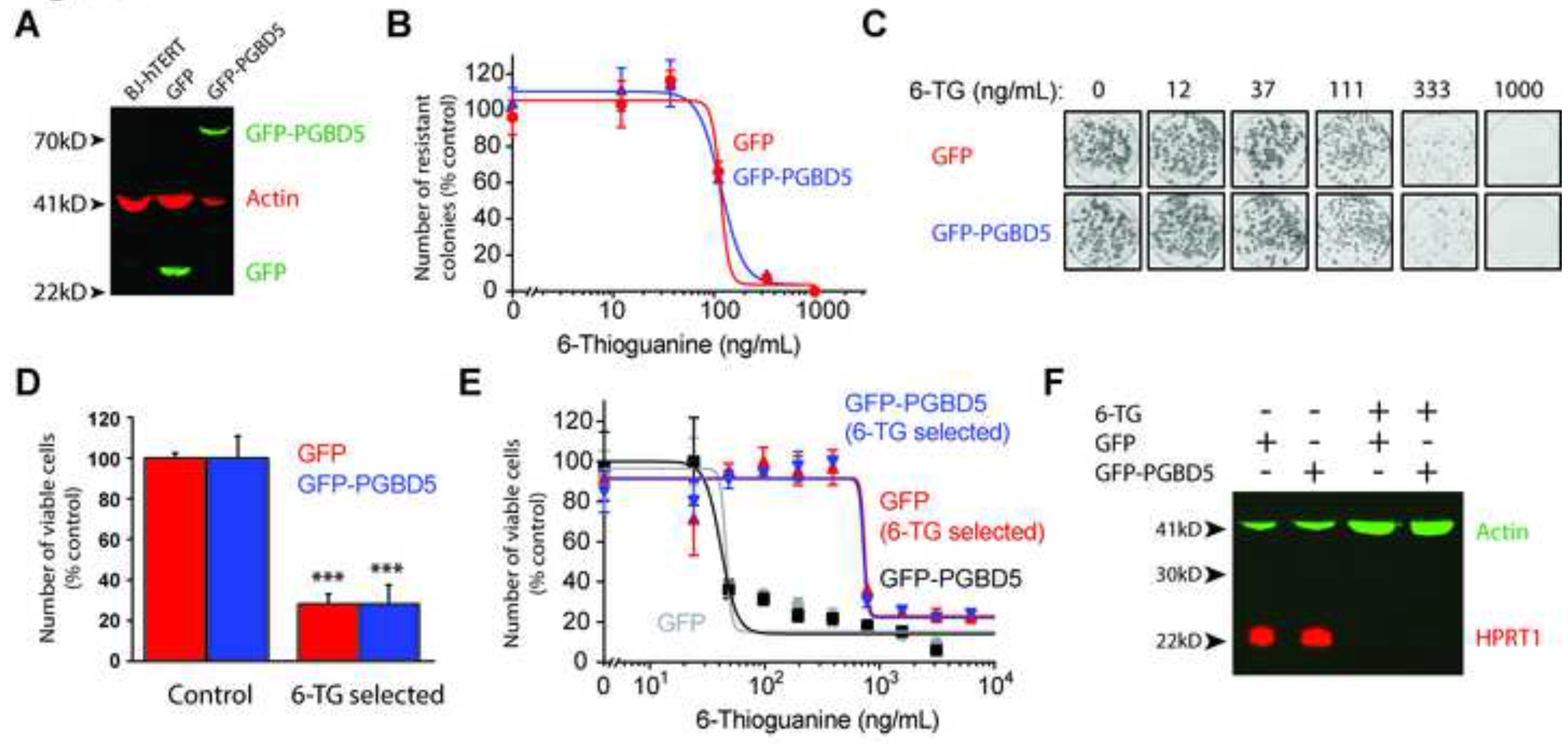
Induction of thioguanine resistance and loss of HPRT1 expression in isogenic cells expressing DNA transposase PGBD5. (**A**) Stable expression of GFP-PGBD5 and control GFP in BJ-hTERT cells, as assessed by Western blotting against GFP; P-actin serves as loading control. (**B**) Clonogenic efficiency of BJ-hTERT cells stably expressing GFP (red) and GFP-PGBD5 (blue) as a function of varying thioguanine concentrations upon thioguanine resistance selection. (**C**) Representative photographs of resistant colonies stained with Crystal Violet. (**D**) Thioguanine selection of both GFP and GFP-PGBD5 expressing cells yields cells that lack HPRT1 activity, as assessed by hypoxanthine-aminopterin-thymidine (HAT) treatment; ^***^ denotes *p* = 2.2×10^−5^ and 9.6×10^−4^ for the comparisons between control and thioguanine selected GFP and GFP-PGBD5, respectively. (**D**) Thioguanine selection yields thioguanine-resistant cells, as assessed by cellular ATP luminescence assay of GFP (red) and GFP-PGBD5 (blue) expressing cells, as compared to control cells (gray and black). (**E**) Western blot for HPRT1 in BJ-hTERT cells expressing GFP and GFP-PGBD5 upon thioguanine selection; β-actin serves as loading control. All error bars represent standard deviations of 3 biological replicates.

Prior structural studies of *HPRT1* gene mutations have used Southern blotting or cDNA sequencing (19), limiting the detection and sequence analysis of structural variants that involve rearrangements of introns and other non-exonic sequences. To overcome these limitations and enable comprehensive and sensitive sequence analysis of rearrangements involving potential transposon sequences in *HPRT1*, we designed a specific panel of polymerase chain reaction (PCR) amplicons spanning the entire 43 Kb sequence of the human *HPRT1* gene, including introns, exons, and 5’ and 3’ untranslated regions (Fig. 2A and Supplemental Table 2). Massively parallel paired-end DNA sequencing of the resultant amplicons, producing millions of sequence reads spanning the *HPRT1* locus, should enable the recovery of rare DNA mutations without the need for single cell cloning that is limited by clonal fitness, as observed for RAG1-induced *HPRT1* mutations in lymphocytes (20, 21).

**Figure 2.**
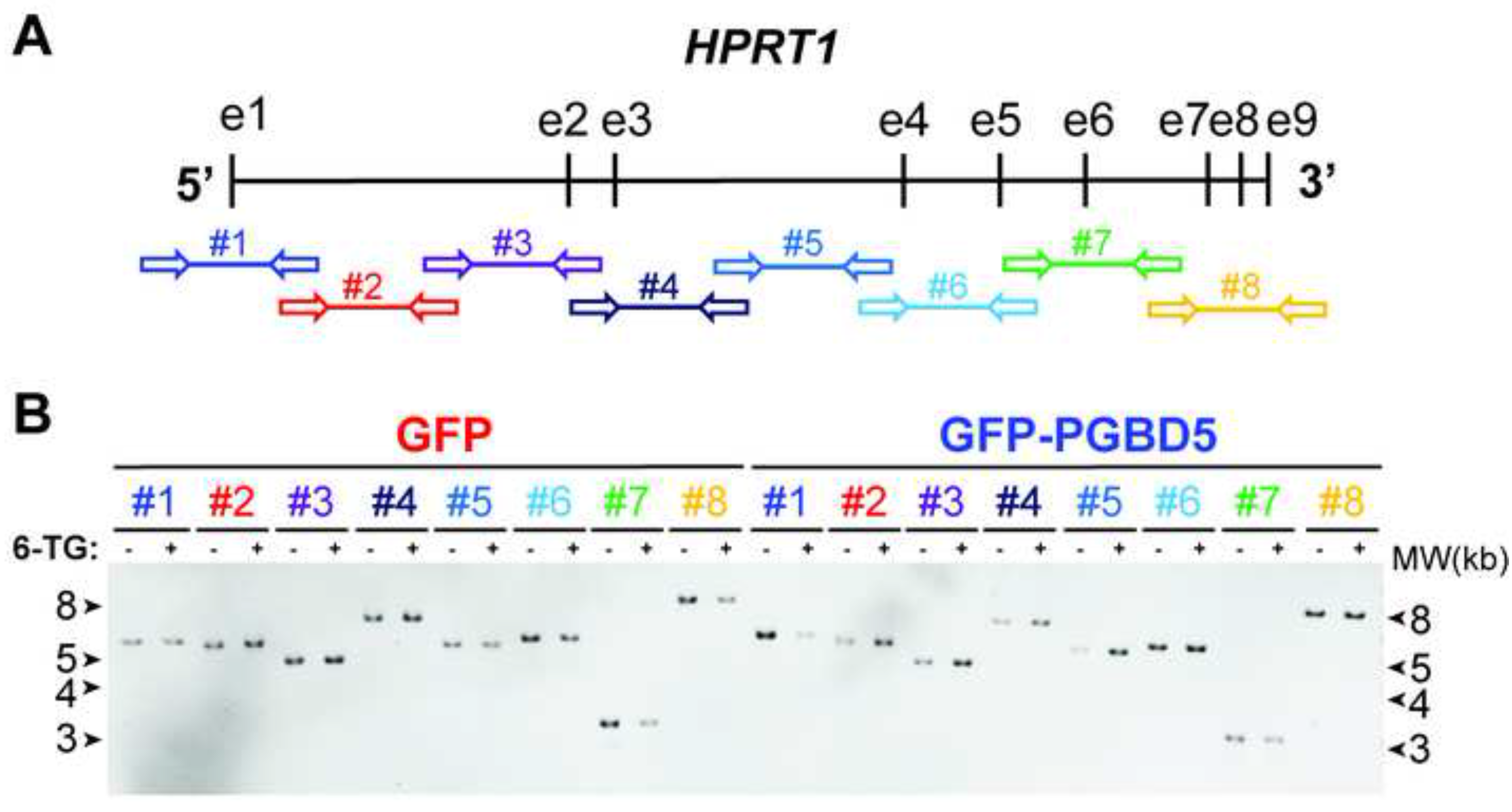
Massively parallel DNA sequencing combined with PCR amplification for high-resolution mutational analysis of *HPRT1*. (**A**) Schematic of the human *HPRT1* gene structure with vertical bars and horizontal arrows denoting exons and amplicons as numbered. (**B**) Photograph of ethidium bromide-stained and electrophoretically-resolved *HPRT1* PCR amplicons of genomic DNA isolated from GFP (red) and GFP-PGBD5 (blue) expressing cells before (−) and after (+) thioguanine resistance selection.

To test this prediction, we isolated genomic DNA from thioguanine-resistant cells expressing GFP-PGBD5 or GFP, and amplified their *HPRT1* loci using long-range PCR (Fig. 2A). Consistent with prior observations of *HPRT1* mutations that were either subclonal or involved variants not resolvable by electrophoresis (14, 15, 21), resultant amplicons exhibited no apparent differences in electrophoretic gel mobility between thioguanine-resistant and control cells in the presence or absence of PGBD5 (Fig. 2B). To facilitate the recovery of polyclonal populations of *HPRT1* mutants, we used massively parallel paired-end Illumina DNA sequencing of resultant genomic amplicons to generate more than 32,000 sequence reads at 99% of nucleotide bases. These data have been deposited to the Sequence Read Archive (http://www.ncbi.nlm.nih.gov/sra/ accession number SRP068848), with the processed and annotated data available from the Dryad Digital Repository (http://dx.doi.org/10.5061/dryad.t748p).

To enable comprehensive analysis of PGBD5-induced *HPRT1* mutations, we combined two recently developed algorithms CRISPResso and laSV that permit the identification and analysis of both small and large structural variants in ultra-high coverage DNA sequencing data at base-pair resolution (22, 23). We observed that both GFP and GFP-PGBD5 expressing cells acquired single nucleotide and small indel mutations of *HPRT1* (Fig. 3A). Consistent with the functional loss of HPRT1 expression (Fig. 1F), we observed a relative excess of exonic as compared to intronic variants in thioguanine-resistant cells (Fig. 3A, right). There was no significant difference in the frequencies of single nucleotide and small indel mutations of *HPRT1* between GFP and GFP-PGBD5 expressing cells (Fig. 3A, left). Although both GFP-PGBD5 and GFP expressing cells developed thioguanine resistance at least in part due to the acquisition of inactivating single nucleotide and small indel mutations, this finding suggests that PGBD5 activity on human genomic substrates does not directly generate such mutations at the efficiency required for thioguanine resistance selection.

**Figure 3.**
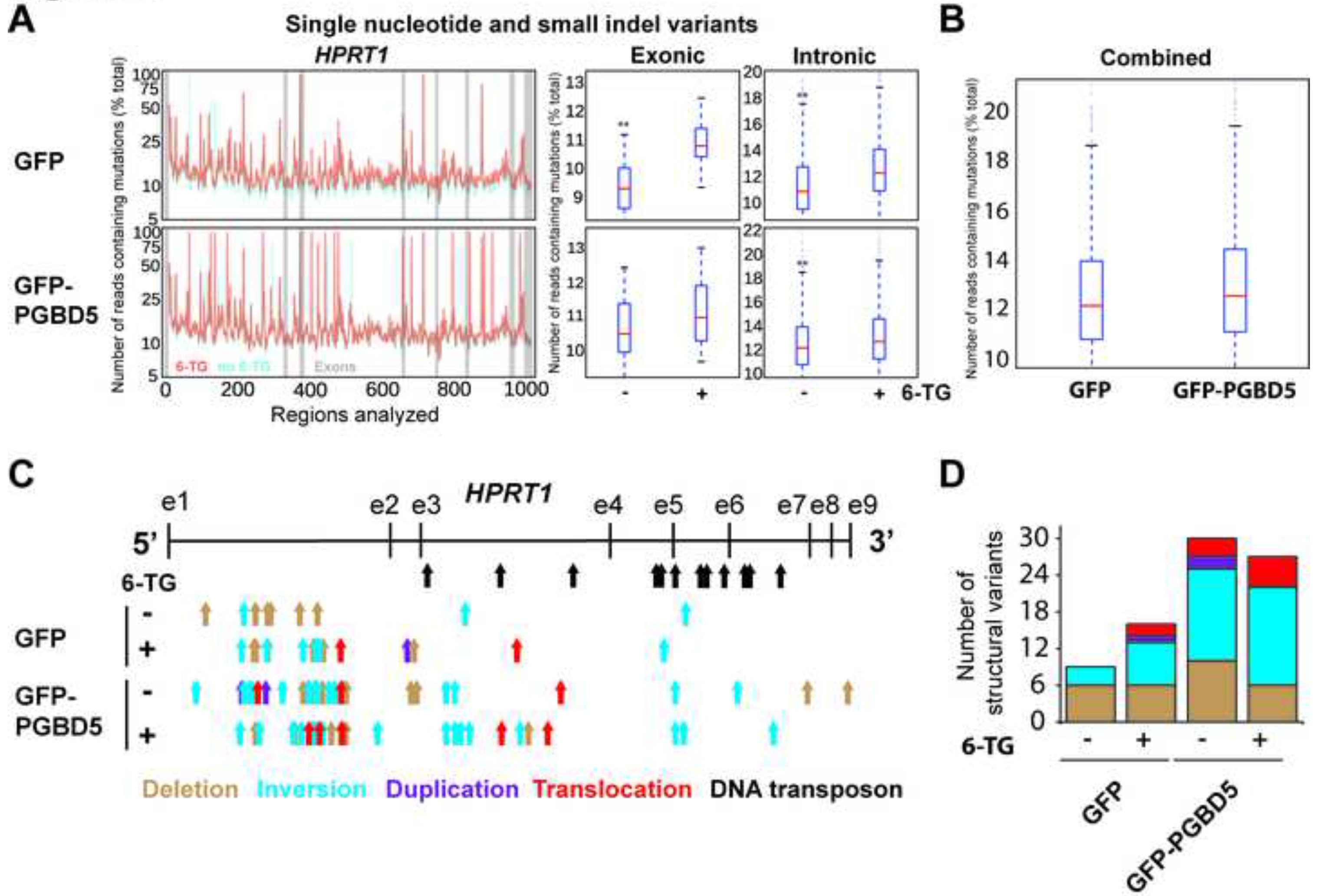
Comprehensive genomic analysis of *HPRT1* mutations reveals PGBD5-mediated induction of complex genomic rearrangements. (**A**) (left) Distribution of the mutational frequency (y-axis) and the location (x-axis) of single nucleotide variants (SNV) and small indels in *HPRT1* of cells before (blue) and after (orange) thioguanine resistance selection. Exons are denoted by gray bars. (right) Comparative analysis of the frequencies of SNVs and indels in *HPRT1* before (−) and after (+) thioguanine resistance selection reveals no significant differences between GFP and GFP-PGBD5 expressing cells; * and ** denote *p* < 0.05 and *p* < 0.01 for exonic and intronic variants, respectively (Exonic GFP *p* = 8.32e-6, exonic GFP-PGBD5 *p* = 0.09, intronic GFP *p* = 2.77e-36, intronic GFP-PGBD5 *p* = 4.77e-06). (**B**) Combined comparative analysis of the frequencies of SNVs and indels in *HPRT1* in GFP and GFP-PGBD5 expressing cells *(p* = 7.90e-4). (**C**) Distribution of the locations of the 5’ ends of complex structural variants in cells before (−) and after (+) thioguanine resistance selection, as detected by laSV and marked by arrows denoting deletions (brown), inversions (blue), duplications (purple), and translocations (red). Black arrows mark annotated DNA transposons. (**D**) Expression of GFP-PGBD5 leads to induction of complex structural variants before (−) and after (+) thioguanine resistance selection (Total number of SVs GFP vs. PGBD5, *p* = 0.001, Poisson test, 95% confidence interval 0.262-0.713).

On the other hand, we observed that cells expressing GFP-PGBD5 had an excess of complex structural variants as compared to GFP control cells (Fig. 3C, Fig 3D and Table 1). Specifically, we found that PGBD5-expressing cells contained significantly greater numbers of inversions in *HPRT1* as compared to control GFP cells (*p* = 0.001, Poisson test with an exact reference distribution, 95% confidence interval 0.14-0.68) and also contained significantly greater total numbers of complex structural variants (*p* < 0.001, Poisson test with an exact reference distribution, 95% confidence interval 0.26-0.71) (Fig. 3C). Furthermore we observed that some rearrangements occurred in the absence of thioguanine selection. This is consistent with the preserved nuclease activity of PGBD5 which may induce such structural rearrangements in human genomes by virtue of DNA double strand breaks alone, similar to the RAG1 recombinase (14, 15). PGBD5-induced genomic rearrangements in *HPRT1* did not appear to involve annotated human transposable elements (Fig. 3C), consistent with the absence of canonical *piggyBac*-derived transposons within the human *HPRT1* gene (4).

**Table 1.** Lasv detects significantly more inactivating mutations in thioguanine-resistant cells.

|  | Control | Thioguanine |
| --- | --- | --- |
| **GFP** | 0/10 | 7/17 |
| **GFP-PGBD5** | 12/36 | 10/29 |

Values denote the number of inactivating / total variants detected.

Previously, we found that the genomic integration of transposons by PGBD5 required specific DNA substrate sequences containing inverted terminal repeats with GGG terminal motifs (4). We reasoned that potential human endogenous PGBD5 substrates in *HPRT1* may be identified by the presence of inverted terminal repeats at the sequences flanking the breakpoints of the observed structural variants (specific definition described in the Methods). Using this approach, we identified 13 terminal elements flanking the structural variant breakpoints in the PGBD5-expressing but not in the control GFP cells (Fig. 4 and Table 2). The identified structural variant breakpoint sequences exhibited specific motifs in the PGBD5 but not in the control GFP cells (Fig. 4), as assessed using sequence entropy analysis and implemented in the MEME algorithm (24, 25). Consistent with their thioguanine resistance selection, 2 out of 13 structural variants involving these sequence motifs were predicted to cause inactivation of *HPRT1* by causing exonic deletions, similar to prior studies of RAG1-induced *HPRT1* inactivation (12). We confirmed that the identified PGBD5-specific sequence motifs occur in the *HPRT1* gene at approximately the same frequency as the rest of the human genome, and therefore are not enriched in our assays simply because of their genomic distribution (*p* = 0.066, binomial-test). While they differ in sequence from the canonical *piggyBac*-derived inverted terminal repeats, PGBD5-specific sequence motifs in *HPRT1* are enriched for terminal GGG nucleotides and lack thymines (Fig. 4), in agreement with the structural requirements observed in previous studies using synthetic transposon substrates (4). While these sequences motifs may constitute specific PGBD5 signal sequences (PSS) that are associated with PGBD5-induced genomic rearrangements, it remains to be determined whether PGBD5 induces a similar variety of genomic rearrangements as part of its endogenous activities, or alternatively, whether it mediates true DNA transposition with excision and insertion of specific PSS-containing inverted terminal repeat mobile elements.

**Figure 4.**
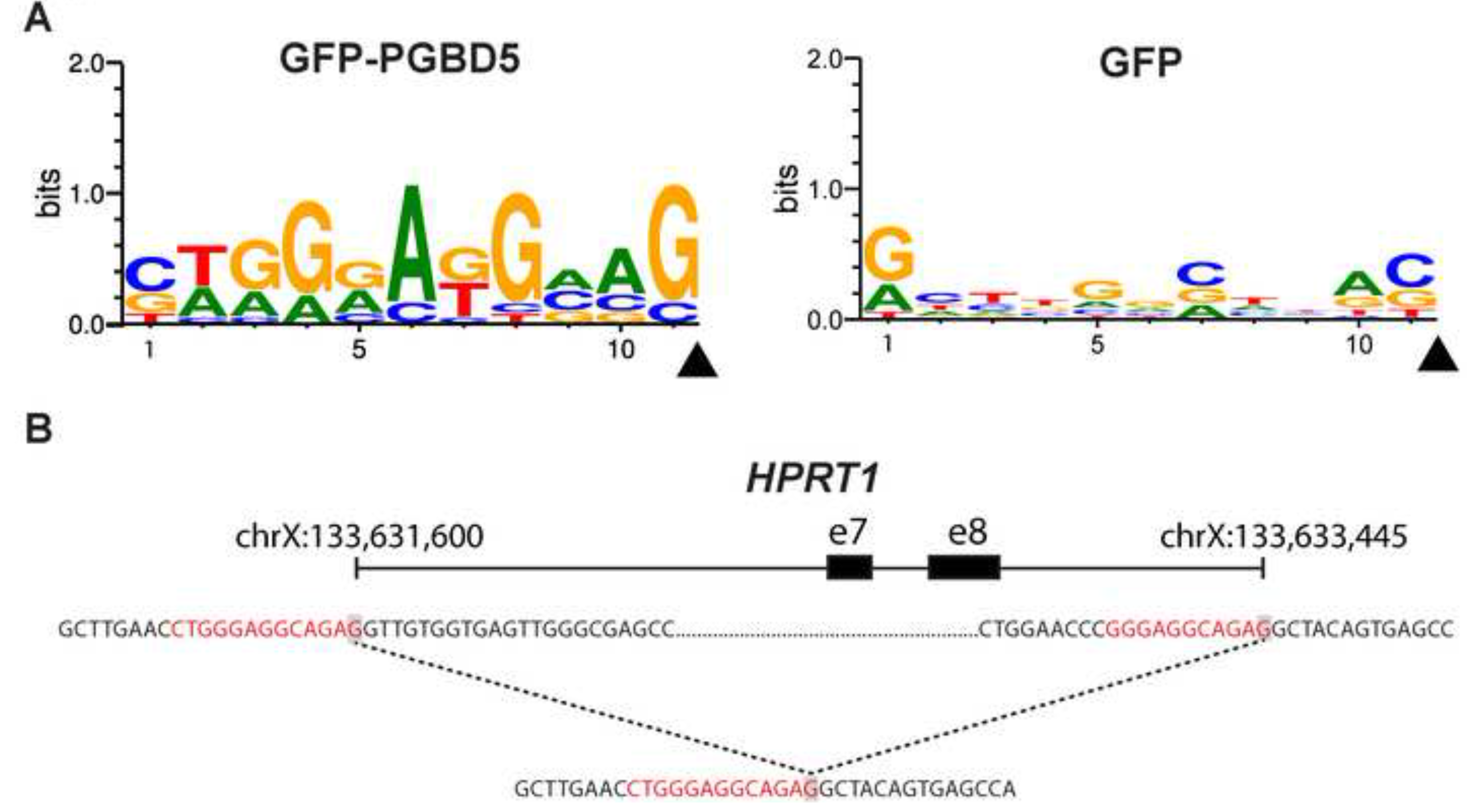
PGBD5-associated *HPRT1* structural variants contain terminal signal sequences. (**A**) Analysis of the sequences flanking structural variant breakpoints demonstrating association of specific signal sequence motifs in cells expressing GFP-PGBD5 (top), but not GFP control (bottom). X-axis denotes nucleotide sequence logo position, and y-axis denotes information content in bits. Black arrowheads mark the location of the breakpoints. (**B**) Breakpoint sequences of a representative structural variant (deletion bnd_11) showing PSS sequence at both breakpoints of a deletion.

**Table 2.**
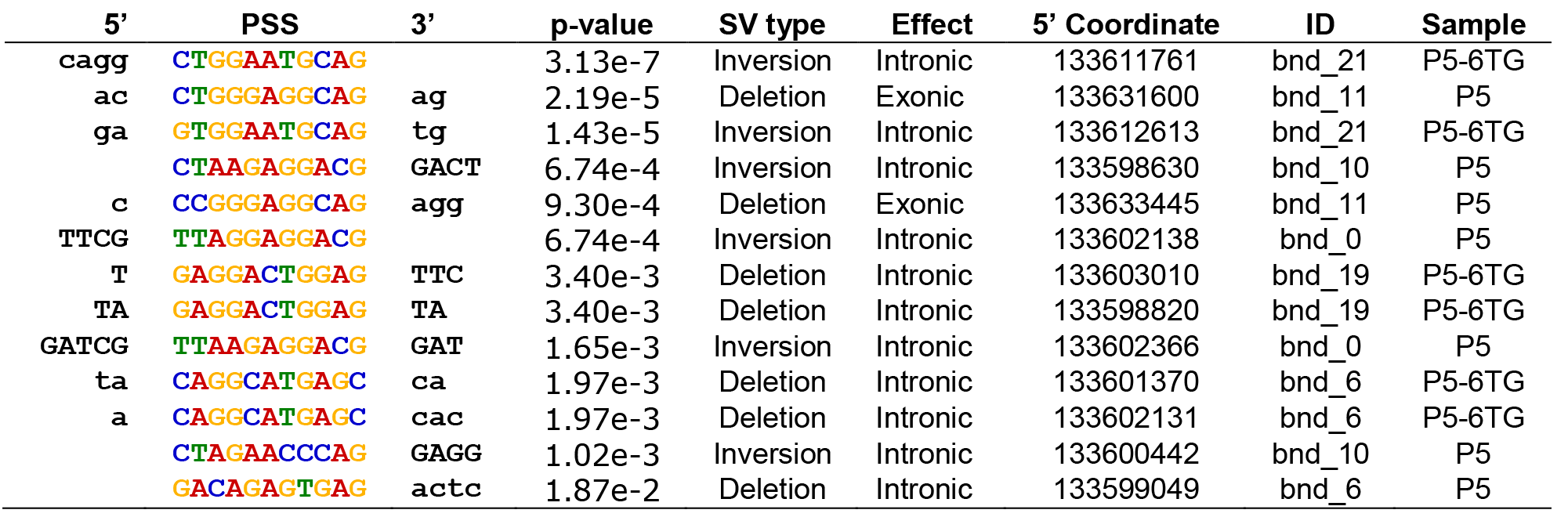
A putative PGBD5 signal sequence (PSS) can be found at breakpoints of structural variants in PGBD5 expressing cells.

Sequences are listed in 5’ to 3’ direction with the breakpoint being on the 3’ end.

## Discussion

In all, our findings indicate that human PGBD5 can induce structural variation and genomic rearrangements of endogenous human *HPRT1* loci. The identification of potential PGBD5 signal sequences in human genomes using the HPRT1 forward genetic screen represents a crucial first step in defining its endogenous genomic substrates in vertebrates and humans. Consistent with the distinct evolutionary history and developmental neuronal expression of PGBD5, identified PSS motifs are distinct from the recombination signal sequences (RSS) described for RAG1 in lymphocytes (14, 15). Importantly, identified PSS motifs exhibit only limited similarity to canonical *piggyBac* transposons, namely preference for terminal GGG nucleotides, in support of the distinct phylogeny of *PGBD5* as compared to other *piggyBac*-derived genes in vertebrates (4). Since our analysis was limited to genomic rearrangements of human *HPRT1* in BJ fibroblasts the presence of thioguanine selection, it is possible that PGBD5 may exhibit different sequence preferences and remodeling activities in neurons and diseased cells where it is endogenously expressed. Our analysis did not identify *bona fide* ‘cut-and-paste’ DNA transposition in *HPRT1*, and it remains to be determined whether PGBD5 catalyzes DNA transposition of endogenous human mobile elements, or simply their nuclease-mediated DNA rearrangements. The described PSS motifs now provide essential templates for future functional studies of PGBD5-induced genomic remodeling.

## Conclusions

Recent discovery of active human THAP9 and PGBD5 DNA transposases, combined with the functional recombination activity of RAG1, suggests that other endogenous transposase-derived genes may catalyze as of yet unknown cell-specific somatic or germ-line rearrangements in vertebrates and humans. While their identification has been substantially empowered by whole-genome sequencing, determination of their functional activities has been hindered by the lack of knowledge of their endogenous substrate sequences. We expect that the integration of forward genetic screening with massively parallel DNA sequencing, as we have done here, and structural variant genome analysis using methods such as CRISPResso/laSV should permit the determination of the genome remodeling activities of endogenous as well as engineered genome editing enzymes. While leveraging the advantages of negative and positive selection of HPRT1 forward genetic screening for specificity, this approach additionally benefits from improved sensitivity, enabling the identification of both simple and complex structural variants at base-pair resolution. This is limited only by sequencing coverage, without the need for single cell cloning that may be compromised by cell fitness effects. Finally, we anticipate that the reported PGBD5 signal sequences will lead to the elucidation of its function in health and disease.

## Methods

### Reagents

All reagents were obtained from Sigma-Aldrich, unless otherwise specified. Synthetic oligonucleotides were synthesized and purified by High Performance Liquid Chromatography (HPLC) by Eurofins MWG Operon (Huntsville, AL, USA).

### Cell culture

BJ-hTERT cells were obtained from the American Type Culture Collection (ATCC, Manassas, Virginia, USA). The identity of all cell lines was verified by Short tandem repeat analysis (STR) analysis and lack of *Mycoplasma* contamination was confirmed by Genetica DNA Laboratories (Burlington, NC, USA). Cell lines were cultured in Dulbecco's Modified Eagle Medium (DMEM) supplemented with 10% fetal bovine serum and 100 U / ml penicillin and 100 μg / ml streptomycin in a humidified atmosphere at 37 °C and 5% CO_2_.

### Plasmid constructs

Human PGBD5 cDNA (Refseq ID: NM_024554.3) was cloned as a GFP fusion into the lentiviral vector pReceiver-Lv103-E3156 (GeneCopoeia, Rockville, MD, USA). Lentivirus packaging vectors psPAX2 and pMD2.G were obtained from Addgene (26). Plasmids were verified by restriction endonuclease mapping and Sanger sequencing, and deposited in Addgene (https://www.addgene.org/Alex_Kentsis/).

### Lentivirus production and transduction

Lentivirus production was carried out as described previously (27). Briefly, HEK293T cells were transfected using TransIT with 2:1:1 ratio of the pRecLV103 lentiviral vector, and psPAX2 and pMD2.G packaging plasmids, according to manufacturer’s instructions (TransIT-LT1, Mirus, Madison, WI). Virus supernatant was collected at 48 and 72 hours post-transfection, pooled, filtered and stored at −80 °C. BJ-hTERT cells were transduced with virus particles at a multiplicity of infection of 5 in the presence of 8 μg/ml hexadimethrine bromide. Transduced cells were selected for 2 days with puromycin (5 μg/ml).

### Western blotting

To analyze protein expression by Western immunoblotting, 1 million transduced cells were suspended in 340 μl of lysis buffer (4% sodium dodecyl sulfate, 7% glycerol, 1.25% beta-mercaptoethanol, 0.2 mg/ml Bromophenol Blue, 30 mM Tris-HCl, pH 6.8). Lysates were cleared by centrifugation at 16,000 *g* for 10 minutes at 4 °C. Clarified lysates (30 μl) were resolved using sodium dodecyl sulfate-polyacrylamide gel electrophoresis, and electroeluted using the Immobilon FLPVDF membranes (Millipore, Billerica, MA, USA). Membranes were blocked using the Odyssey Blocking buffer (Li-Cor), and blotted using antibodies against GFP (mouse anti-human, 1:500, clone 4B10, Cell Signaling Technology, Beverly, MA), β-actin (rabbit antihuman, 1:5000, clone 13E5, Cell Signaling Technology, Beverly, MA), HPRT1 (rabbit antihuman, 1:1000, clone ab10479, Abcam, Cambridge, MA) and β-actin (mouse anti-human, 1:5000, clone 8H10D10, Cell Signaling Technology, Beverly, MA). Blotted membranes were visualized using goat secondary antibodies conjugated to IRDye 800CW or IRDye 680RD and the Odyssey CLx fluorescence scanner, according to manufacturer’s instructions (Li-Cor,Lincoln, Nebraska).

### Hypoxanthine-aminopterin-thymidine (HAT) medium selection

HAT medium prepared using the 50X HAT supplement (Thermo Fisher Scientific) and DMEM medium with 10% fetal bovine serum and 100 U / ml penicillin and 100 μg / ml streptomycin. Media was replaced twice weekly and cells were grown in the presence of HAT selection for 15 doublings, corresponding to approximately 5 weeks.

### Thioguanine selection

Cells were cultured in the presence of 120 ng/ml of 6-thioguanine for 10 doublings, corresponding to approximately 4 weeks. Media was replaced twice weekly.

### Cell viability and colony formation assays

For cell viability assays, cells were seeded at a density of 200,000 cells per well in 6-well plates (Corning Life Sciences, Corning, NY, USA). Twenty four hours after seeding, medium was replaced with HAT medium. The number of viable cells was counted 2 days after treatment using Trypan Blue staining using the Neubauer hematocytometer according to the manufacturer’s instructions (Thermo Fisher Scientific).

For clonogenic assays, cells were seeded at a density of 10,000 cells per 10-cm dish and treated with 6-thioguanine (0-1 μg/ml) for 2 weeks. Resultant colonies were fixed with methanol, stained with Crystal Violet, and counted manually using a spatial grid.

To assess cell viability of cells after treatment with 6-thioguanine, cells were seeded into 9-well plates (Thermo Fisher Scientific, Waltham, MA, USA) at a density of 1,000 cells per well. Cell were treated with 6-thioguanine (0-1 μg/ml) twenty four hours after seeding. Cell viability was quantified using the CellTiter-Glo ATP content luminescence based assay, according to manufacturer’s instructions (Promega, Madison, WI, USA).

### Generation of *HPRT1* amplicons

Genomic DNA was extracted from 10 million cells using the PureLink Genomic DNA Mini Kit according to the manufacturer’s instructions (Thermo Fisher Scientific). To exponentially amplify the *HPRT1* gene, we designed primer pairs every 3-8 Kb (Suppl. Table 2). Amplicons were generated using 50 ng of gDNA in 50 μl reaction volumes containing 0.5 μM of primers. Loci 2 to 8 where amplified using the Phusion Green High-Fidelity DNA Polymerase (Thermo Scientific, Waltham, MA, USA) with the following parameters: 98°C for 30 sec, followed by 30 cycles of 98 °C for 10 sec, 65 °C for 30 sec and 72 °C for 3.5 min, and a final extension of 72 °C for 3.5 min. Locus 1 was amplified using the KAPA Long Range HotStart DNA Polymerase (KAPA Biosystems, Wilmington, MA, USA) with the following parameters: 94 °C for 3 min, followed by 40 cycles of 94 °C for 25 sec, 60 °C for 15 sec and 72 °C for 7 min, and a final extension of 72 °C for 7 min. PCR products were purified using the PureLink PCR purification kit according to the manufacturer’s instructions (Invitrogen Corp., Carlsbad, CA, USA).

### Illumina library preparation and sequencing

Equimolar amounts of purified PCR amplicons were pooled, as measured using fluorometry with the Qubit instrument (Invitrogen Carlsbad, CA) and sized using the BioAnalyzer 2100 instrument (Agilent Technologies, Santa Clara, CA). The sequencing library construction was performed using the KAPA Hyper Prep Kit (KAPA Biosystems, Wilmington, MA) and 12 indexed Illumina adaptors obtained from IDT (Coralville, IO), according to the manufacturer’s instructions. After quantification and sizing, libraries were pooled for sequencing on a MiSeq (pooled library input at 10 pM) using a 300/300 paired-end run (Illumina, San Diego, CA). A total of 728,000-928,000 paired reads were generated per sample. The duplication rate varied between 0.22 and 0.27%. The data reported in this manuscript have been deposited to the Sequence Read Archive (SRA) (http://www.ncbi.nlm.nih.gov/sra/, accession number SRP068848), and the Dryad Digital Repository (http://dx.doi.org/10.5061/dryad.t748p).

### Mutational and structural variant analysis

For the analysis of single nucleotide and small indel mutations, we used the CRISPResso WGS utility from the CRISPResso software using default parameters (22). The analysis was performed on non-overlapping windows of 40bp spanning the entire *HPRT1* gene body. For the analysis of large structural variants, we used laSV with the following parameters: −s 30 −k 63 −p 30 (23).

### PGBD5 signal sequence analysis

Clustal Omega with default parameters was used for multiple sequence alignment (24). PGBD5 signal sequences were defined using the following criteria: i) sequences flanking 5’ and 3’ breakpoints demonstrated at least 50% identity less than 4 bp from the breakpoints when aligned to each other in inverted orientation, ii) aligned sequences contained no single or tandem repeats longer than 5 bp, and iii) no such alignments were identified at breakpoints of variants found in GFP control expressing cells. Sequence motifs were identified using MEME with default parameters by referencing alignments in the 5’ to 3’ direction with the breakpoint at the 3’ terminus (28).

### Statistical analysis

Mutational frequencies were calculated as described previously (18), according to the following formula: Mutational frequency = −Ln(*X_s_* / *N_s_*) / −Ln(*X_0_* / *N_0_*), where *N* is the number of cells seeded and *X* is the number of colonies formed with (S) and without (0) thioguanine selection. The difference between the number of mutations across samples were compared using a Poisson test with an exact reference distribution.

## Availability of data and material

Data deposition: The data reported in this manuscript have been deposited to the Sequence Read Archive (SRA) (http://www.ncbi.nlm.nih.gov/sra/, accession number SRP068848), the Dryad Digital Repository (http://dx.doi.org/10.5061/dryad.t748p).

## Ethics and consent

Ethics approval was not required for any aspect of this study.

## Abbreviations

PSS: PGBD5-specific signal sequences
HAT: hypoxanthine-aminopterin-thymidine
RSS: recombination signal sequences
PCR: polymerase chain reaction
SV: structural variant
SNV: single nucleotide variant
STR: Short tandem repeat analysis
HPLC: High Performance Liquid Chromatography
DMEM: Dulbecco’s Modified Eagle Medium
6-TG: 6-thioguanine

## Author contributions

A.H. and A.K. designed the research, A.H., E.J., C.V. performed the research, A.H., L.P., J.Z., Z.W., D.B., A.K. analyzed data, A.H. and A.K. wrote the manuscript in consultation with all authors.

## Acknowledgments

We thank David Pellman for suggesting forward chemical genetic screening for transposon discovery and Cedric Feschotte for helpful comments on the manuscript.

## Supplementary files

**Supplementary Table 1.** Annotated DNA transposons in the *HPRT1* gene.

| Position | Transposon Name | Transposon Class |
| --- | --- | --- |
| ChrX: 133609504-133609757 | MARNA | TcMar-Mariner |
| ChrX: 133612862-133613034 | MER104 | TcMar-Tc2 |
| ChrX: 133616475-133616651 | MER3 | hAT-Charlie |
| ChrX: 133623197-133623402 | Charlie7a | hAT-Charlie |
| ChrX: 133623432-133623581 | Eulor8 | TcMar |
| ChrX: 133624726-133624872 | MER5A | hAT-Charlie |
| ChrX: 133625413-133625487 | Tigger5 | TcMar-Tigger |
| ChrX: 133625475-133625553 | MER47C | TcMar-Tigger |
| ChrX: 133626916-133626974 | MER5A | hAT-Charlie |
| ChrX: 133628672-133628708 | MADE1 | TcMar-Mariner |
| ChrX: 133628926-133629162 | MER106B | hAT-Charlie |
| ChrX: 133630729-133630913 | MER5B | hAT-Charlie |

**Supplementary Table 2.**
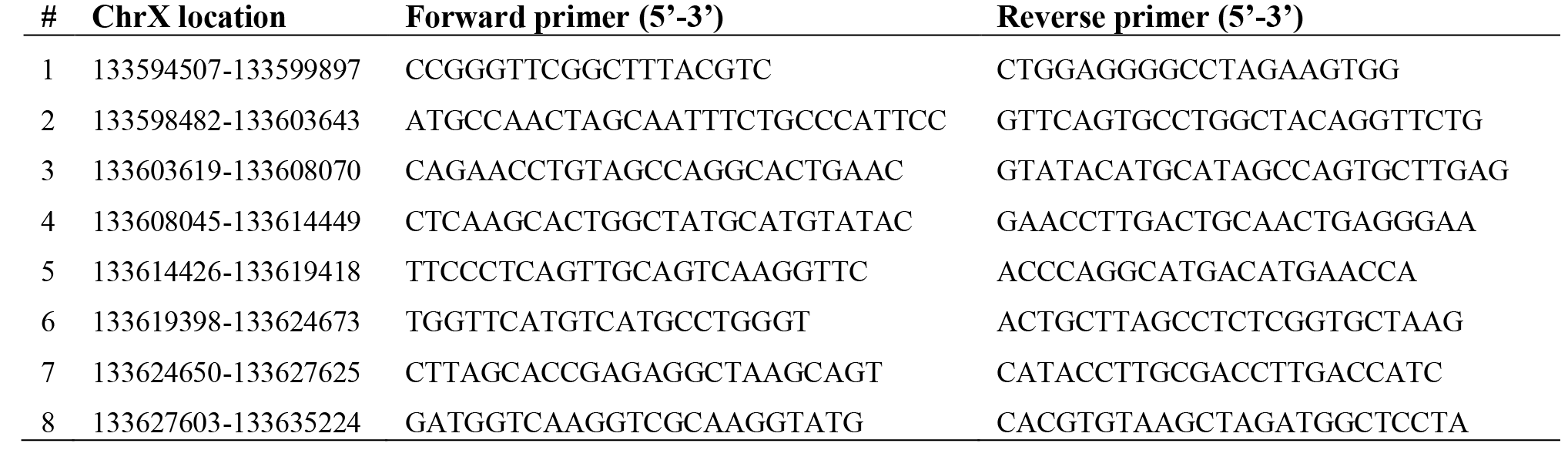
HPRT1 PCR primers and PCR amplicon locations.

